# A positive association between gut microbiota diversity and vertebrate host performance in a field experiment

**DOI:** 10.1101/2023.07.20.549938

**Authors:** Andreas Härer, Ken A. Thompson, Dolph Schluter, Diana J. Rennison

## Abstract

The vertebrate gut microbiota is a critical determinant of organismal function, yet it remains unclear if and how gut microbial communities affect host fitness under natural conditions. Here, we investigate associations between growth rate (a fitness proxy) and gut microbiota diversity and composition in a field experiment with threespine stickleback fish (*Gasterosteus aculeatus*). We detected on average 63% more bacterial taxa in the guts of high-fitness fish compared to low-fitness fish (i.e., higher α-diversity), suggesting that higher diversity promotes host growth. The microbial communities of high-fitness fish had higher similarity (i.e., lower β-diversity) than low-fitness fish, supporting the Anna Karenina principle— that there are fewer ways to have a functional microbiota than a dysfunctional microbiota. Our findings provide a basis for functional tests of the fitness consequences of host-microbiota interactions.

**Significance statement:** The vertebrate gut microbiota is important for many aspects of their hosts’ biology—such as nutrient metabolism and defense against pathogens—that could ultimately affect host fitness. However, studies investigating the effects of gut microbiota composition on vertebrate host fitness under natural conditions remain exceedingly rare. We tested for associations between gut microbiota diversity and growth rate (a fitness proxy) in threespine stickleback fish reared in large outdoor ponds. We found evidence that a more diverse gut microbiota was predictive of higher growth rate and therefore increased host fitness. Notably, high-fitness fish had higher gut microbiota similarity to one another than did low-fitness fish, providing experimental evidence for the Anna Karenina principle—that there are fewer ways to have a functional microbiota than a dysfunctional microbiota—as it relates to host fitness.

## Introduction

Host-microbe interactions are a universal feature of eukaryotes, and a combination of environmental and host factors (e.g., physiology, morphology, and ecology) shape gut microbiota diversity (1, 2). Several lines of evidence indicate that microbiota have important effects on host fitness. For example, microbes can facilitate host adaptation to novel environments by enabling metabolism of dietary resources, and variation in ecologically relevant phenotypes can be mediated by microbes (3, 4). However, most insights into fitness effects of host-microbiota interactions come from invertebrate model systems reared under laboratory conditions (5). Because many environmental variables in the laboratory are substantially different from natural environments, it is an important endeavor to evaluate host fitness-gut microbiota relationships in natural settings to better understand the impact on host evolutionary trajectories (6). Yet, making inferences in wild individuals can be complicated because host fitness is influenced by a range of abiotic and biotic factors that are often difficult to account for or measure. By controlling external factors, experiments conducted under natural conditions provide a powerful opportunity to determine the effects of the gut microbiota on host fitness.

To this end, we leveraged a unique field-based experimental infrastructure to investigate whether gut microbiota diversity is associated with differences in growth rate (a fitness proxy) in threespine stickleback fish (*Gasterosteus aculeatus*; hereafter ‘stickleback’). We introduced different combinations of ecologically divergent benthic and limnetic ecotypes originating from three lakes in British Columbia, Canada, into three experimental ponds and characterized the gut microbiota of individuals with the lowest (low-fitness fish) and highest (high-fitness fish) growth rates from each source population and pond (Fig. 1A). Stickleback are well-suited for studying gut microbiota-mediated fitness effects because body size is positively correlated with fitness via higher fecundity and overwinter survival (7). We predicted a positive association between gut microbiota α-diversity and host performance, since higher α-diversity has been suggested to be beneficial for the host (8, 9). Next, we predicted higher similarity in gut microbiota composition (i.e., lower β-diversity) among high-fitness fish, which follows from the Anna Karenina principle that has been argued to be common in animal microbiomes (10). Our study provides novel insights by assessing host fitness-gut microbiota associations on the individual host level in free-living animals under semi-natural conditions.

**Fig. 1:**
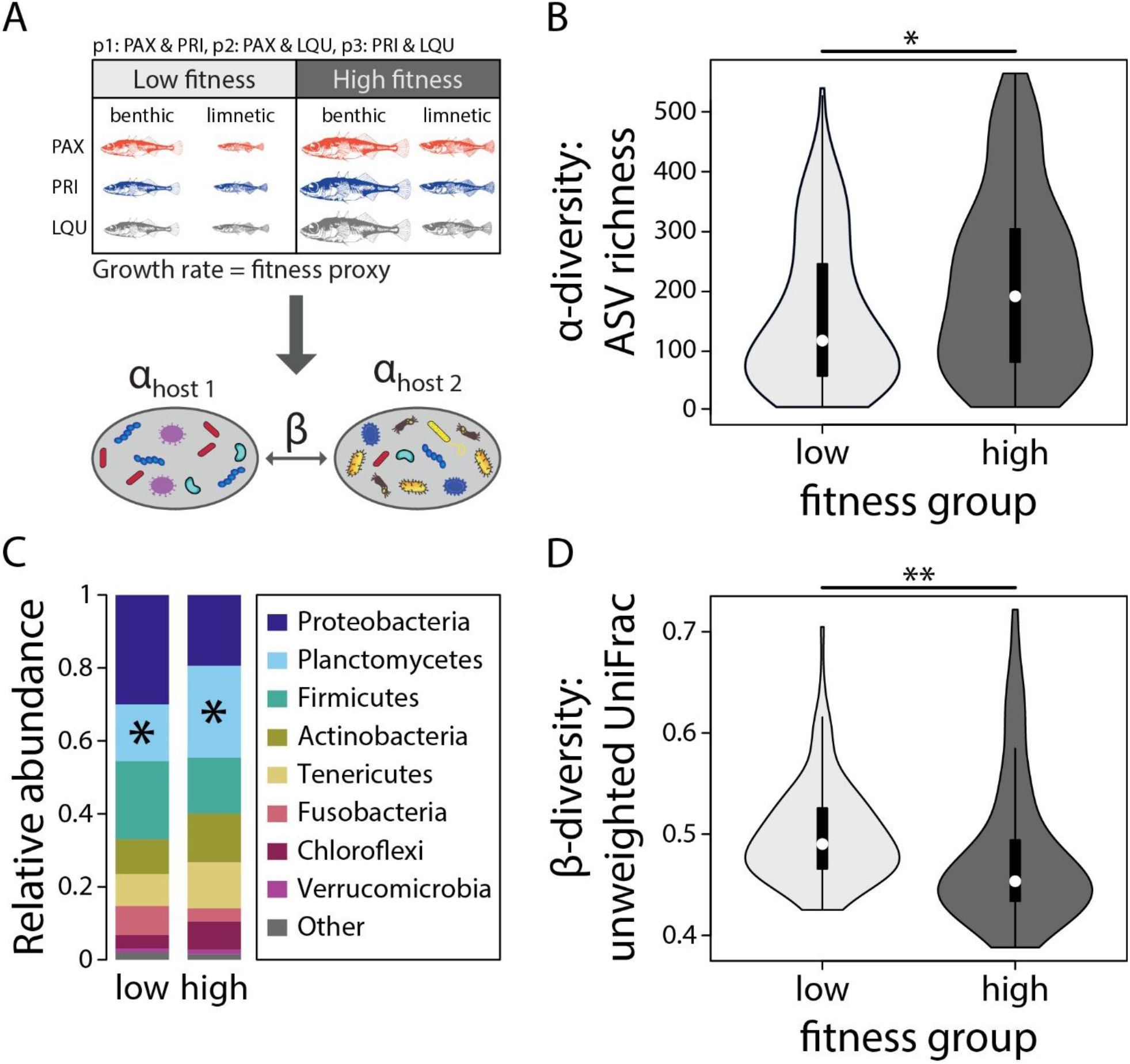
Lab-raised benthic and limnetic fish from Paxton Lake (PAX), Priest Lake (PRI), and Little Quarry (LQU) were introduced into three ponds in different combinations; their fitness was assessed by growth rate and gut microbiota α- and β-diversity was determined (A). High-fitness fish had higher α-diversity (B), higher abundance of Planctomycetes (C), and lower β-diversity (distance from fitness group centroid) (D), which together provides evidence for substantial gut microbiota differences between fitness groups (n_low-fitness_ = 121, n_high-fitness_ = 119). \**P* < 0.05, \*\**P* < 0.01.

## Results

The bacterial diversity associated with single hosts (α-diversity) differed significantly between fitness groups for amplicon sequence variant (ASV) richness (linear mixed-effects model; *F*_1,204.24_ = 10.60, *P* = 0.001), Faith’s phylogenetic diversity (*F*_1,209.71_ = 7.76, *P* = 0.006), and Shannon diversity (*F*_1,220_ = 6.77, *P* = 0.010) (Fig. 1B). Median ASV richness (192 vs 118) and Faith’s phylogenetic diversity (18.78 vs 13.98) were 63% and 34% higher in high-fitness fish compared to low-fitness fish, but median Shannon diversity was only 4% higher in high-fitness fish (3.85 vs 3.69). Within the low-fitness group, relative fitness values were positively correlated with ASV richness (Spearman’s rank correlation; *ρ* = 0.20, *P* = 0.025) and Faith’s phylogenetic (*ρ* = 0.26, *P* = 0.004) but no correlations were apparent within the high-fitness group (all *P* > 0.05). The cumulative ASV richness (γ-diversity) was similar for the low- and high-fitness groups (4,343 and 4,387 ASVs; *P* = 0.186); 2,374 ASVs were present in both fitness groups whereas 1,969 and 2,013 ASVs were only found in the low- and high-fitness group, respectively. While no ASVs were shared among all members of a fitness group, the major bacterial phyla were similar across fitness groups and only Planctomycetes were more abundant in high-fitness fish (ANCOM; *W* = 22) (Fig. 1C).

We found evidence for significant gut microbiota dissimilarity (β-diversity) between low- and high-fitness fish (PERMANOVA; unweighted UniFrac: *F*_1,228_ = 3.07, *P* = 0.001), a pattern that was likely driven by lower β-diversity within the high-fitness group than within the low-fitness group (PERMDISP; unweighted UniFrac: *F*_1,239_ = 6.96, *P* = 0.004) (Fig. 1D). For inferred metagenome function, we found no evidence for dissimilarity between fitness groups (MetaCyc pathway abundance: *F*_1,228_ = 0.85, *P* = 0.463), but again there was lower β-diversity within the high-fitness group (KEGG Orthologs: *F*_1,239_ = 9.65, *P* = 0.002). Only one MetaCyc pathway (sucrose degradation III) was differentially abundant between fitness groups with a higher abundance in low-fitness fish. Overall, these results indicate that gut microbial communities, and their inferred metagenomic function, are more similar among high-fitness fish than among low-fitness fish.

## Discussion

Our study represents an important first step in establishing a connection between individual host fitness and gut microbiota composition in a free-living vertebrate, highlighting that variation in microbial communities can have important consequences for their hosts (3).

Why is microbial α-diversity positively associated with individual growth rate? No bacterial ASV was shared among all members of the high-fitness group, and we found little evidence for differentially abundant bacterial phyla or metabolic pathways, which in combination indicates that faster growth is unlikely to be driven by a few beneficial taxa. Further, both fitness groups harbor a similarly diverse—and partially overlapping—pool of bacteria with equal proportions of unique taxa. However, high-fitness fish, on average, capture a larger proportion of this γ-diversity within each host individual. While the exact mechanisms by which higher α-diversity might promote host growth remain to be determined, our results suggest that a more diverse gut microbiota per se could be beneficial, for example, by allowing hosts to metabolize a broader range of nutrients and/or exploit novel trophic niches (11).

High- and low-fitness groups had significantly dissimilar gut microbiota (β-diversity), and our study is the first to demonstrate such an association under semi-natural conditions in a vertebrate host. This pattern appeared to be driven by higher similarity in gut microbiota composition and inferred metagenome function (i.e., lower β-diversity) within the high-fitness group (Fig. 1D). In line with the Anna Karenina principle (10), our results suggest that ‘all microbiota increasing host fitness are similar; each microbiota decreasing host fitness does so in its own way’. Yet, which exact microbiota configurations might promote higher host fitness remains to be explored. Since gut microbes can shape a range of host phenotypes and life history, variation in gut microbiota composition could further impact other fitness-related traits (3, 6, 12, 13).

Importantly, we overcame limitations of previous studies by measuring gut microbiota composition and a fitness proxy in individual hosts reared under semi-natural conditions. This setting standardized both the early-life diet and the age of fish, and allowed fish to occupy and forage in different microhabitats; dietary patterns in our experimental ponds (as evidenced by variation in isotope values) correspond to those observed in wild stickleback populations (14). This is crucial because the effects of gut microbiota variation on host fitness can depend on environmental factors (especially diet) (15), emphasizing that the ecological context may strongly affect host-microbiota interactions.

In summary, we demonstrate that variation in gut microbial communities is associated with a proxy for stickleback fitness, but it remains to be determined whether the observed variation is a cause or consequence of differences in host fitness. Hence, our work provides a foundation for future work aiming to establish causal relationships through gut microbiota manipulation (13). We advocate that such studies be conducted in a diverse range of host organisms to determine the generality of host fitness-gut microbiota interactions.

## Methods

Fish were weighed and implanted with numbered tags, then released into experimental ponds for 71–99 days. Growth rate was assessed for surviving individuals via the change in body mass while controlling for initial body mass and number of days in the experimental ponds. For each source population and pond, we selected 8–12 individuals per fitness group for gut microbiota analysis based on amplicon sequencing of the 16S rRNA gene (n_low-fitness_ = 121, n_high-fitness_ = 119).

To determine if gut microbiota diversity impacts fitness, we tested for differences in bacterial gut microbiota α-diversity, which measures the bacterial diversity of individual host fish (linear mixed-effects models; ASV richness, Faith’s phylogenetic diversity, and Shannon diversity). To test the Anna Karenina principle, we measured β-diversity between individual fish, which captures the (dis)similarity of bacterial communities (PERMANOVA, PERMDISP; Bray-Curtis dissimilarity, unweighted UniFrac, weighted UniFrac) and inferred metagenome function (MetaCyc pathway abundance) between hosts. Analyses included the effects of host ecotype, sex (determined genetically), genetic background (lake-of-origin), and diet (carbon and nitrogen isotope signatures) as fixed effects and rearing environment (pond) as a random effect. Results were consistent across β-diversity metrics, and we only report unweighted UniFrac statistics in the main text. Statistics for the other two metrics, as well as detailed methods for library preparation and statistical analyses including effects of aforementioned factors on gut microbiota diversity, in addition to potentially confounding artifacts due to sequencing (e.g., read number), are provided in the Supporting Information. Differential abundance of bacterial phyla and MetaCyc pathways was assessed by analysis of composition of microbiomes (ANCOM).

## Acknowledgments

This work was supported by funding from the Deutsche Forschungsgemeinschaft (DFG, German Research Foundation) – project number 458274593 to AH and from the University of California San Diego to DJR. KAT was supported by a Human Frontier Science Program Long-Term Fellowship. DS was supported by the Natural Sciences and Engineering Research Council of Canada. D. Rubenstein alerted us to the Anna Karenina principle.

## Supporting Information for

## Experimental design

All fish used for this study were part of an experiment studying the fitness consequences of hybridization (no hybrids were used herein), and we refer to Thompson & Schluter (1) for more detailed information. Parents of the experimental fish were raised from hatching in a common laboratory environment and pure within-population crosses were made between unrelated fish of benthic and limnetic populations (variously called ‘species’ or ‘ecotypes’) from each of Paxton Lake, Priest Lake, and Little Quarry Lake, in British Columbia, Canada. At present, these are the only three lakes with extant benthic-limnetic species pairs; it is possible that more await discovery (2). Their offspring were raised in aquaria until the juvenile stage and fed a common diet. For the experiment, fish were kept in unmanipulated semi-natural ponds (25 × 15 m including benthic and limnetic habitats) at the University of British Columbia, Canada (3) and all fish were approximately the same age at the start of the experiment. In each pond, there were combinations of benthic and limnetic populations from two lakes each (Fig. 1A), and fish were kept in the ponds between 71 to 99 days. Fish were weighed before and after the experiment, and sequential coded wire tags (Northwest Marine Technology, Anacortes, WA, USA) were used for individual identification. There were no weight differences between low-fitness and high-fitness fish at the start of the experiment (two sample t-test, *t* = 0.264, *P* = 0.792).

## Data collection

Fish were collected using minnow traps or dip nets and were then immediately euthanized with an overdose of MS-222, after which they were weighed, photographed, and stored at − 20°C in individually labelled 15mL tubes. Growth rate for each individual fish—our fitness proxy—was calculated based on mass gain during the experiment with linear models using final mass as the response variable and initial weight and the days an individual spent in the pond during the experiment as fixed effects and pond as a random effect. Models for benthic and limnetic ecotypes were fitted separately due to substantial differences in body size and allometry, see Thompson & Schluter for more details (1). In total, we sampled approximately 20 fish per population within each pond; 10–12 fish with low relative fitness and 10–12 fish with high relative fitness. Some samples were later excluded due to low sequencing depths, and final sample sizes ranged from 8–12 fish per group which accumulated to a total of 121 and 119 low- and high-fitness fish.

Before beginning our sequencing library preparation, all fish were rinsed with 95% EtOH and their whole guts were dissected using sterile equipment. We subsequently carefully removed any gut contents then stored samples stored at − 80 °C. DNA was extracted from whole guts with the QIAGEN PowerSoil Pro Kit according to the manufacturer’s protocol (Qiagen, Hilden, Germany) under sterile conditions in a laminar flow hood. To characterize microbial communities associated with fish guts, we amplified the V4 region of the 16S rRNA gene with barcoded 515F and 806R primers (https://github.com/SchlossLab/MiSeq_WetLab_SOP/blob/master/MiSeq_WetLab_SOP.md). All PCR reactions were done in triplicate using the Q5 High-Fidelity 2X Master Mix (New England Biolabs, Ipswich, MA) and pooled for each fish after amplification. The PCR had a denaturation step for 60 s at 98 °C, 35 amplification cycles with 10 s at 98 °C, 20 s at 56 °C and 60 s at 72 °C, and a final elongation at 72 °C for 10 min. To check for successful amplification, we visualized PCR products by gel electrophoresis (2% agarose gel), and DNA concentrations were measured on a Qubit 4 Fluorometer (Thermo Fisher Scientific, Waltham, MA). We included negative controls of sterile water for DNA extraction and PCRs, none of which yielded any detectable DNA amplification. All samples were subsequently pooled in an equimolar manner for the two libraries (samples were either sequenced in 2021 or 2022). At the UC Davis Genome Center, libraries were purified by bead clean-up and DNA quality was checked on a Bioanalzyer. The final libraries were sequenced on the Illumina MiSeq 600 (PE300) platform.

To obtain information on diet, we collected muscle tissue to determine stable isotope ratios of carbon (δ^13^C) and nitrogen (δ^15^N), which allow detecting diet variation associated with benthic and limnetic habitats in stickleback (3, 4). Muscle tissues were dried at 55°C and subsequently homogenized to a powder, 1 mg of each sample was loaded into a tin capsule and combusted in a Elementar vario EL cube elemental analyzer interfaced to an Elementar VisION IRMS (Elementar Analysensysteme GmbH, Germany) at the UC Davis Stable Isotope Facility. Laboratory standards indicated measurement errors (SD) of ± 0.05‰ for δ^13^C and 0.07‰ for δ^15^N. We further collected fin tissue of each fish in 95% EtOH, and after DNA extraction the sex of each fish was determined by PCR following the protocol develop by Peichel et al. (5).

## Data analysis

Our data consisted of a total of 9,493,085 raw sequencing reads (mean: 39,554 reads/sample). For some samples, we obtained low sequencing depths, which were further decreased by merging of forward and reverse reads due to filtering during this step. Hence, we chose to use 250 bp of the forward reads for our gut microbiota analyses, since these reads consistently showed higher sequence quality compared to reverse reads yet encompassed 86% of the target locus (250 out of 291 bp). All upstream analyses described below were done in QIIME2 (6). In order to obtain amplicon sequencing variants (ASVs), sequencing reads were checked for quality and corrected, and chimeric sequences were removed using the *dada2* plugin (7). Next, FastTree 2.1.3 was used to assemble a phylogenetic tree of the bacterial lineages (8), and bacterial taxonomy was assigned based on the SILVA 132 ribosomal RNA (rRNA) database at a 99% similarity threshold (9). Before conducting downstream analyses, we excluded ASVs with less than 10 reads that were detected only in a single sample and ASVs that could not be assigned at the class level. We further filtered out ASVs mapping to chloroplasts, mitochondria, cyanobacteria, or archaea to obtain information of the bacterial gut microbiota only. ASV counts were normalized through scaling with ranked subsampling (SRS) with a C_min_ of 2500 reads (10). To infer metagenome function, MetaCyc pathway abundances were predicted with the PICRUSt2 plugin in QIIME2 (11) with a maximum nearest-sequenced taxon index (NSTI) cutoff of 2.

First, we tested for effects of ecotype, lake-of-origin, carbon and nitrogen stable isotope, sex, fitness group, and pond on α-diversity (ASV richness, Faith’s phylogenetic diversity, Shannon diversity) and β-diversity (non-phylogenetic: Bray-Curtis dissimilarity, phylogenetic: unweighted and weighted UniFrac; 12, 13). For α-diversity, we used linear mixed effect-models (*lmer* function in lme4 package v1.1-31) (14) with pond as random effect and all other variables as fixed effects and produced analysis-of-variance tables to test for statistical significance of the model terms (*anova* function in stats package v4.2.1) (15). Besides fitness group, we detected an effect of host sex on all three α-diversity metrics (ASV richness: *F*_1,219.39_ = 7.70, *P* = 0.006, Faith’s phylogenetic diversity: *F*_1,218.96_ = 6.08, *P* = 0.014, Shannon diversity: *F*_1,220_ = 7.93, *P*= 0.005), and an effect of carbon signature on Shannon diversity (*F*_1,220_ = 5.74, *P* = 0.017). Adding sequencing read number as another independent variable to the models showed that it had a significant effect on all three α-diversity metrics, but difference in α-diversity between fitness groups were maintained, indicating that the observed effects were not driven by differences in read numbers. Moreover, all differences in α-diversity were evident when examining rarefaction plots that illustrate α-diversity as a function of sequencing depth for both fitness groups. We further used Spearman’s rank correlation to test for correlations between fitness values and α-diversity within each fitness group. For comparing gut microbiota dissimilarity within and between fitness groups (β-diversity), we used PERMANOVA (*adonis2* function in vegan package v2.6-2) (16, 17), and we found that pond, sex, and lake-of-origin had significant effects on dissimilarity of gut microbiota taxonomic composition across all three metrics. Further, host ecotype, as well as carbon and nitrogen signatures showed significant effects based on Bray-Curtis dissimilarity and weighted UniFrac. Notably, gut microbiota were significantly dissimilar between fitness groups after statistically controlling for the effects of all other factors. For inferred metagenome function, we found significant effects of ecotype, lake-of-origin and nitrogen signature on β-diversity. We then determined the cumulative ASV richness for each fitness group (γ-diversity). To test for significant differences in γ-diversity between fitness groups, we resampled individual hosts with replacement 10,000 times for each fitness group, calculated γ-diversity for each iteration and determined the γ-diversity ratio between the high-fitness group and the low-fitness group. We then tested for higher γ-diversity in the high-fitness group by calculating the proportion of iterations for which the high-fitness group had a higher γ-diversity and we determined statistical significance using a cut-off of 0.05.

To test whether the magnitude of gut microbiota dissimilarity measured within each fitness group differs between fitness groups, we determined β-diversity dispersion by calculating the distance of each fish from the centroid of its respective fitness group (*betadisper* function in vegan package v2.6-2). We used the *adonis2* function (vegan package v2.6-2) to calculate *P*-values for the comparison of β-diversity values between fitness groups. For gut microbiota taxonomic composition, we found that β-diversity was lower for high-fitness fish for all three metrics: Bray-Curtis dissimilarity (F_1,239_ = 7.32, *P* = 0.007), unweighted UniFrac (F_1,239_ = 6.96, *P* = 0.004), and weighted UniFrac (F_1,239_ = 5.02, *P* = 0.031). Since conclusions drawn about host fitness from the analysis of β-diversity metrics were consistent across metrics, we decided to only report the unweighted UniFrac metric in the main text. All statistical analyses were done in R v4.2.1 (18).

## Notes

### Competing Interest Statement

The authors have declared no competing interest.

